# EphrinA4/EphA4 controls blood pressure via arterial sympathetic innervation

**DOI:** 10.1101/2023.02.03.526852

**Authors:** Emilie Simonnet, Sabrina Martin, José Vilar, Emilie Vessieres, Sonia Taib, Virginie Monceau, Luc Pardanaud, Nadine Bouby, Anne Eichmann, Jean-Sébastien Silvestre, Daniel Henrion, Isabelle Brunet

**Affiliations:** Center for Interdisciplinary Research in Biology (CIRB), College de France, CNRS, INSERM, Université PSL, Paris, France.; Université de Paris, PARCC, INSERM, F-75015 Paris, France.; U 1083, Angers, France; Centre de Recherche des Cordeliers, INSERM, Sorbonne Université, Université de Paris, F-75006 Paris, France; Radiotoxicology and Radiobiology Research Laboratory (LRTOX), Institute of radiation and nuclear Safety (IRSN), Fontenay-aux-roses, France

**Author notes:** Equal contribution (Emilie Simonnet, Sabrina Martin).

## Abstract

The autonomic sympathetic nervous system innervates peripheral resistance arteries, thereby controlling arterial diameter and modulating blood supply to organs and arterial tone. Despite its fundamental role in blood flow regulation and adaptive response of the cardiovascular system to challenging situations, how sympathetic arterial innervation develops remains poorly understood.

We here show that sympathetic arterial innervation is regulated by the axonal guidance molecule EphrinA4 in arterial Smooth Muscle Cells (SMCs), which repels sympathetic axons via the EphA4 receptor. Specific inactivation of EphA4 in sympathetic axons induced a loss of repulsion and increased sympathetic innervation of peripheral arteries throughout life. Functional consequences were a significant increase in arterial tone (resistivity and vasoconstriction), leading to an elevated systemic arterial blood pressure that reached to hypertension under stressful circumstances. These findings identify a novel pathway that negatively regulates sympathetic arterial innervation, and could participate to the appearance of idiopathic resistant hypertension.

## Introduction

The sympathetic nervous system innervates internal organs and regulates physiological body functions but also fine-tunes the adaptative response to challenging situations; such as stress or immediate danger. Sympathetic neurons aggregate during development into ganglions to form the sympathetic ganglion chain that lies along the spinal cord (1). Axonal fibers exit cell bodies and extend over long distances to innervate smooth muscle cells in internal organs (2–4). To reach their distant targets, sympathetic axons follow arteries, which produce secreted cues that guide axon extension (2, 5). This occurs in mice around E15,5. Later during development, starting from postnatal day 2 (P2), arteries themselves attract sympathetic axons, and resistance arteries get fully innervated in a lace-like pattern by P10 (6). Neurovascular junctions (NVJs) form between sympathetic fibers and vascular SMC (vSMC) (7) ; those varicosities are “en passant” synapses responsible for neurotransmitter release. Noradrenaline release fosters vSMC contraction and subsequently arteriole constriction, thereby controlling vascular tone and participating to blood pressure regulation (8).

Netrin-1and VEGF have been shown to regulate the onset of arterial innervation guidance and NVJs patterning (6, 9). Netrin-1 is guiding sympathetic axons toward developing arteries via the neuronal receptor Deleted in Colorectal Cancer (DCC) and is involved in the maintenance of sympathetic innervation and NVJs formation. Genetic inactivation of Netrin-1 resulted in a decreased arterial innervation and reduced NVJ number and size. Remarkably, inactivation of Netrin-1 in adult animals was sufficient to reduce innervation, suggesting that arterial innervation is a dynamic and finely regulated process. Of note, sympathetic axons have the ability to regenerate (10, 11), entailing that sympathetic innervation level can vary over time depending on axonal signaling perceived. At a functional level, the rate of sympathetic innervation in arteries can be involved in tissue homeostasis and pathological conditions (3, 12). In the context of essential hypertension, renal artery denervation is still considered as a potential treatment for resistant hypertension (13). In this line of reasoning, spontaneous hypertensive rats have hyper-innervated arteries, and sympathectomy normalizes their blood pressure levels (14). Finally, during pregnancy, preeclampsia is characterized by a marked increase in peripheral vascular resistance, which reverts to normal after delivery. Such an increase in blood pressure is mediated, at least in part, by a substantial activation of sympathetic vasoconstriction (15).

This prompted us to asked whether sympathetic innervation of arteries was a regulated and refined process, during lifetime. We reasoned that the level of innervation could be finely controlled by a balance of attractive and repulsive cues adapting and maintaining arterial innervation rate.

## Results

### EphrinA4/EphA4 trigger sympathetic axon repulsion from arteries

We identified in newly innervated postnatal mesenteric arteries using whole-mount ISH the expression of EfnA4, which encodes the repulsive axon guidance molecule EphrinA4. EfnA4 is expressed at the innervation onset by mesenteric arteries, at Postnatal day 2 (P2) (Figure 1A). EfnA4 is expressed by arterial smooth muscle cells (SMC) as EfnA4 mRNA was visualized by Fluorescent In Situ Hybridization in acta2 positive cells of the arterial SMC layer at P2 (Figure 1B). EphrinA4 is expressed at the membrane of arterial SMC as early as P2, and persists at P15 in primary arterial vSMC cultured until day 7 *in vitro* (Figure 1C). EphA4, a protein-tyrosine kinases receptor for EphrinA4 (16, 17), was detected on sympathetic axons (Figure 1D) and neurons (Figure 1E) from the Superior Cervical Ganglion from P1 to adulthood. To test if EphrinA4 could bind sympathetic neurons, we performed binding experiments using recombinant Fc-tagged EphrinA4 protein incubated with sympathetic neurons isolated from wildtype mice and grown *in vitro*. Anti-Fc labeling showed that EphrinA4 bound to sympathetic axon shafts and growth cones (Figure 1, F and H). Binding was lost in sympathetic neurons from EphA4-deficient (EphA4-/-) mice (18) (Figure 1, G and H and Supplemental Figure 1, A and B), identifying EphA4 as the obligate EfnA4 receptor on sympathetic axons. As EphrinA4 triggers contact-mediated repulsion (19), we tested effects of EphrinA4 on sympathetic axons from wildtype mice *in vitro* using collapse assay which uses the morphology of the growth cone that collapse after exposure to repellent cues (20). We found that EphrinA4 mediated collapse of sympathetic axons a dose-dependent manner (Figure 1I). The collapse response was abolished when sympathetic axons isolated from EphA4-/- mice (Figure 1, J and K), demonstrating that EphrinA4-mediated repulsion required the EphA4 receptor. Hence, arterial SMC expressing EphrinA4 could repel sympathetic axons via the receptor EphA4.

**Figure 1.**
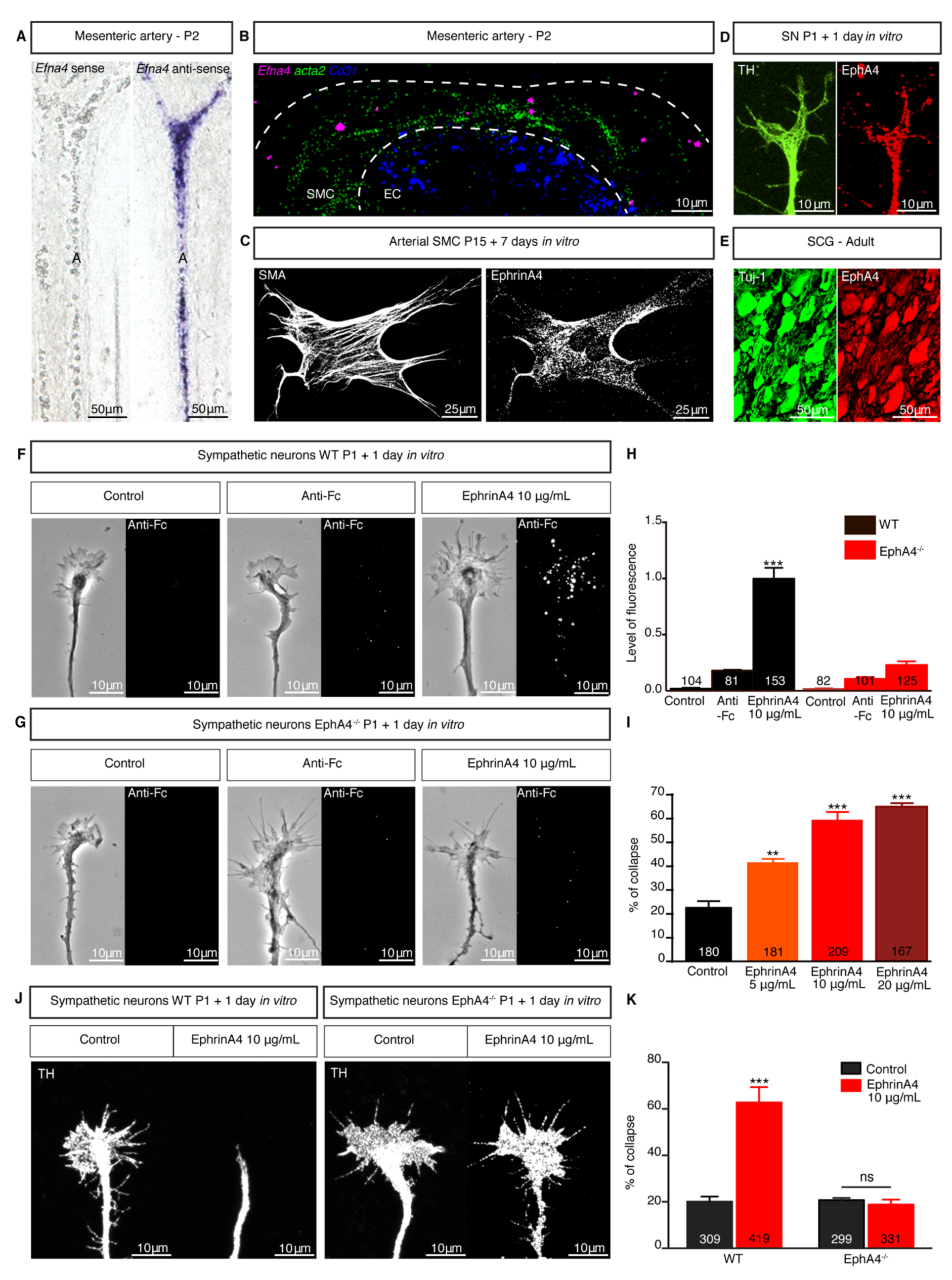
EphrinA4-EphA4 are expressed upon sympathetic arterial innervation and mediate axonal repulsion. (**A**) *In situ* hybridization (ISH) of *Efna4* mRNA in whole-mount mesenteric artery (A) from WT mice at P2. Control *Efna4* sense probe (left) and anti-sense probe (right) ensured staining specificity. (**B**) Fluorescent ISH of *Efna4* (magenta), *acta2* (green) and *cd31* (blue) in transverse sections of mesenteric arteries from WT mice (P2). Doted lines delineate smooth muscle cells (SMC) layer and endothelial cells (EC). (**C-E**) Immunofluorescent staining of SMA (smooth muscle actin, left) and EphrinA4 (right) on an arterial SMC (P15 WT mouse mesentery) cultured *in vitro* (7 days) (C); of TH (Tyrosine hydroxylase, green) and EphA4 (red) on sympathetic neurons (SN) from Superior Cervical Ganglia (SCG) of a WT mouse (P1) after 1 day *in vitro* (**D**); of Tuj-1 (green) and EphA4 (red) in a transverse section of adult SCG from WT mice (E). (**F and G**) Binding assay on SN from WT (F) and EphA4^-/-^ mice (**G**) collected at P1, and cultured *in vitro* during 1 day. SN were stimulated with either control media (left), CY3- anti-Fc (middle) or clustered EphrinA4-Fc-CY3 (10 μg/mL, right). (H) Quantification of fluorescence levels of CY3-Anti-Fc signal on those SN. (**I**) Quantification of axonal collapse percentage of WT SN stimulated with EphrinA4 (from 0 to 20 μg/mL). (**J**) Collapse assay of SN (expressing TH) from WT and EphA4^-/-^ mice at P1, cultured *in vitro* during 1 day, stimulated with control media (left) or clustered-EphrinA4 (10 μg/mL, right). (**K**) Quantification of the collapse assay. ** p<0.01 ***p<0.001

### Loss of EphA4-mediated repulsion leads to increased arterial innervation

To evaluate if EphrinA4 and EphA4 regulated arterial innervation *in vivo*, we used *EfnA4* ^-/-^ and EphA4 ^-/-^ mice (Supplemental Figure 1, A and B) (in fact EfnA4^-/-^ is a triple Knock-out (TKO) resulting in the simultaneous deletion of Efna1Efna3Efna4 located in the same genomic region (21), but only EphrinA4 is expressed in arteries (not shown)). We investigated the onset of arterial innervation in pups at P3. To visualize sympathetic axons, we stained mesenteries with an antibody against tyrosine hydroxylase (TH), the rate-limiting enzyme in catecholamine production in the sympathetic nervous system (8, 22), and to visualize arteries, we used an antibody directed against Smooth Muscle Actin (SMA). We quantified the area of TH+ axons covering the SMA+ arterial wall at P3 and observed a significant increase of arterial innervation in EphrinA4-/- and EphA4 ^-/-^ mice compared to WT littermates (Figure 2A). Arterial diameter was not affected (Figure 2B). We then deleted *EphA4* in sympathetic axons using *EphA4*^flox/flox^ mice and *TH-*Cre driver lines (23, 24) (hereafter designated EphA4^flox^-TH^CRE^). Cre negative littermates were used as controls in this study. While SCG from EphA4^flox^-TH^CRE^ expressed half of the wildtype EphA4 levels (Figure 2, D and E and Supplemental Figure 1C), sympathetic arterial innervation was increased (Figure 2, A-D, Supplemental Figure 1C). Thus, reduction of EphA4 levels by only 50% is sufficient to enhance arterial innervation.

**Figure 2.**
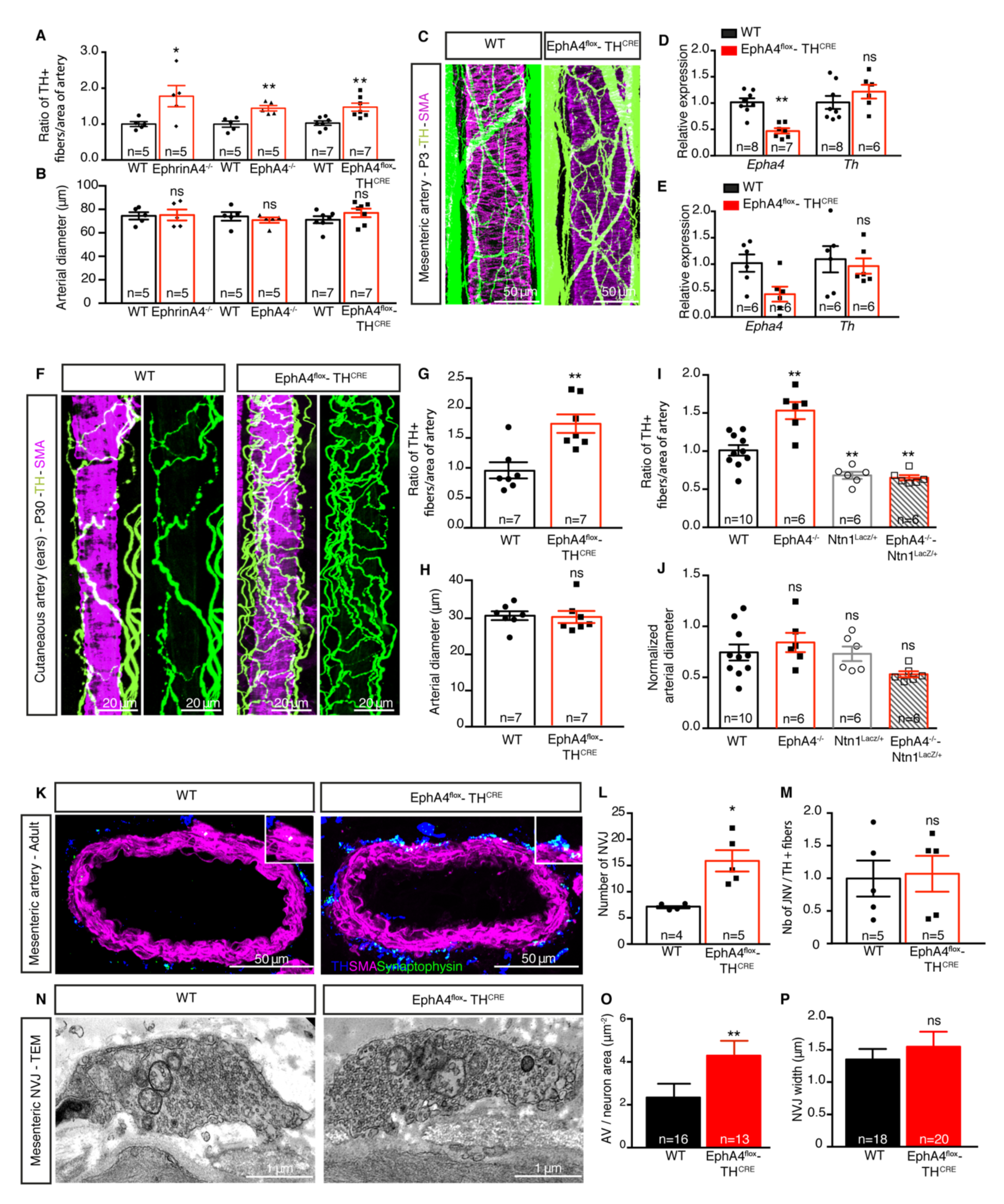
Loss of EphrinA4-EphA4 signaling *in vivo* leads to enhanced sympathetic arterial innervation and NVJs, compatible with a loss of repulsion. (**A and B**) Quantification of percentage of TH+ nerve fibers covering mesenteric arteries (A) and arterial diameter (B) from WT, EphrinA4^-/-^, EphA4^-/-^ and EphA4^flox^- TH^CRE^ mice (P3). (**C**) Whole-mount immunofluorescent staining of TH (green) and SMA (magenta) on mesenteric arteries from WT (left) and EphA4^flox^-TH^CRE^ mice (P3). (**D and E**) Relative normalized expression of *Epha4* and *Th* mRNA in SCG from WT and EphA4^flox^-TH^CRE^ at P3 (D), and from adults (E). (**F**) Immunofluorescent staining of TH (Sympathetic nerves, green) and SMA (SMC, magenta) on cutaneous arteries (ears) from WT (left) and EphA4^flox^-TH^CRE^ (right) mice (P30). (G and H) Quantification of the percentage of TH+ nerve fibers covering cutaneous arteries (G) and quantification of the arterial diameter (H) from WT and EphA4^flox^-TH^CRE^ mice (P30). (I and J) Quantification of TH+ nerve fibers percentage covering cutaneous arteries from WT, EphA4^-/-^, Ntn1^LacZ/+^ and EphA4^-/-^-Ntn1^LacZ/+^ mice (P30) (**I**) and quantification of the arterial diameter normalized to WT animals for each genotype (J). (**K**) Immunofluorescent staining of TH (blue), SMA (magenta) and synaptophysin (green) on mesenteric artery sections from adult WT and EphA4^flox^-TH^CRE^ mice. Boxes show close-up of neurovascular junctions (NVJ). (**L**) Quantification of the number of NVJ on mesenteric arteries from adult WT and EphA4^flox^-TH^CRE^ mice. (**M**) Ratio between number of NVJ and TH+ fibers covering the mesenteric arteries for each genotype, normalized to WT value. (N) Transmission electronic microscopy images of NVJ in mesenteric arteries from adult WT and EphA4^flox^-TH^CRE^ mice. (**O**) Quantification of the number of Adrenergic Vesicles (AV) divided by NVJ area. (**P**) Quantification of NVJ width on mesenteric arteries from adult WT and EphA4^flox^-TH^CRE^ mice. ns: not significant, * p<0.05; ** p<0.01 ;***p<0.001.

To test if such arterial hyperinnervation persisted throughout adulthood, we evaluated arterial innervation in 30 days-old mice. Relative EphA4 expression was still significantly lower in EphA4^flox^-TH^CRE^ animals compared to WT littermates, both at the mRNA and protein levels (Figure 2E and Supplemental Figure 1D). Arterial innervation was significantly enhanced by more than 50% in EphA4^flox^-TH^CRE^ mice compared with control animals while arterial diameter was unaffected (Figure 2, F-H). The number of EphA4+ neurons was decreased in EphA4^flox^-TH^CRE^ SCG, but the total number of neurons per SCG was similar and unaffected both during development and in adult animals (Supplemental Figure 2, A and B). Similarly, arterial innervation was increased in EphA4-/- animals using both in whole mount and cryosection quantification of TH+ fibers density (Supplemental Figure 2, C-H).

As we previously identified Netrin-1 as a positive regulator of arterial innervation (6), we analyzed arterial innervation level when both Netrin-1/DCC and EphrinA4/EphA4 signaling were altered. We crossed EphA4-/- mice with mice inactivated for Netrin-1 (25). As the Netrin-1^LacZ/LacZ^ mice die at birth, we used the Netrin-1 heterozygote mice that were already described as exhibiting a significant decrease of arterial innervation (6). As expected, we found that arterial innervation was significantly increased in EphA4-/- animals compared to WT, whereas a decrease was seen in the Netrin-1^LacZ/+^ animals. Interestingly, EphA4-/- Netrin-1^LacZ/+^ animals showed a similar phenotype as the Netrin-1^LacZ/+^ animals (Figure 2, I and J). This data suggests that Netrin-1 is primary required for chemo-attraction of sympathetic axons toward the artery. Once axons contact the artery, contact-mediated repulsion is occurring via EphA4-EphrinA4 signaling.

In adults, NVJ controls maturation and function of arterial innervation. We immuno-stained mesenteric arteries with TH and synaptophysin, a pre-synaptic marker, to visualize NVJ (Figure 2K upper panel). We found a significant increase of NVJ number in EphA4^flox^-TH^CRE^ when compared to control animals. Normalization of this number with the percentage of innervation per genotype showed no difference, indicating that the increase of NVJ in EphA4^flox^-TH^CRE^ mice is due to a more abundant innervation, whereas the ratio of NVJ per fiber is conserved (Figure 2, L and M). This data indicates that there are more NVJ because there are more fibers but there are not more NVJ per fiber.

We visualized NVJ using Transmission Electron Microscopy (TEM) and observed similar NVJ size, length, synaptic cleft size or mitochondrial composition. Conversely, the number of adrenergic vesicles per NVJ in the EphA4^flox^-TH^CRE^ mice increased compared to control animals (Figure 2, N-P and not shown). In conclusion, loss of EphrinA4-EphA4 signaling leads to an increased arterial innervation density that remains during adulthood and leading to over-numbered NVJ.

### Functional consequences of an increased arterial innervation

To address the physiological consequences of altered sympathetic innervation, we investigated cutaneous vasoconstriction in response to cold using laser Doppler perfusion imaging. We anesthetized mice and imaged cutaneous blood flow of the paw in adult EphA4^flox^-TH^CRE^ and WT littermates. Blood flow is color coded, red indicating a high blood flow, and blue low blood flow. Under anesthesia, body temperature dropped from 37.5°C to 33.5°C. We quantified cutaneous blood flow every degree’s drop. As expected, WT animals showed significant decrease of cutaneous blood flow in response to cold (between 35.5°C and 34.5°C), indicating that vasoconstriction occurs to maintain body core temperature. EphA4^flox^-TH^CRE^ mice displayed a more rapid (between 36.5°C and 35.5°C) and efficient vasoconstriction (Figure 3, A and B left panel) consistent with their increased arterial innervation. Prazosin, an inhibitor of alpha-adrenergic receptors that mediate contraction of the arterial SMC upon release of catecholamines, was then injected to animal prior experiment. Inhibition of alpha-adrenergic receptors that mediate contraction of the arterial SMC upon release of catecholamines using Prazosin treatment abolished vasoconstrictive response in both wildtype and EphA4^flox^-TH^CRE^ mice suggesting that the change in vasoconstriction efficacy observed could be directly linked to the enhanced sympathetic arterial innervation and NVJ numbers observed in EphA4^flox^-TH^CRE^ mice (Figure 3, A and B right panel). Furthermore, the pulsatility and resistivity index was significantly raised in the carotid artery of EphA4^flox^-TH^CRE^ animals compared to controls (Figure 3, C and D), while arterial diameter and wall thickness were unaffected (Supplemental Figure 3, E-H); in line with a possible increased blood flow and vascular tone. To examine potential changes in arterial wall properties we analyzed arterial wall anatomy using histochemistry. Aortic and mesenteric arteries wall thickness as well as the number and size of SMC layers were unaffected (Figure 3E and Supplemental Figure 3, A-D). Molecular composition was then assessed. RNA extracted from adult second order mesenteric arteries revealed no significant changes in genes expression characterizing mural cell contractility pathways such as acta2, Calponin, Desmin, Smoothelin and Smooth muscle protein 22-alpha or SM22a (Figure 3F) between genotypes.

**Figure 3.**
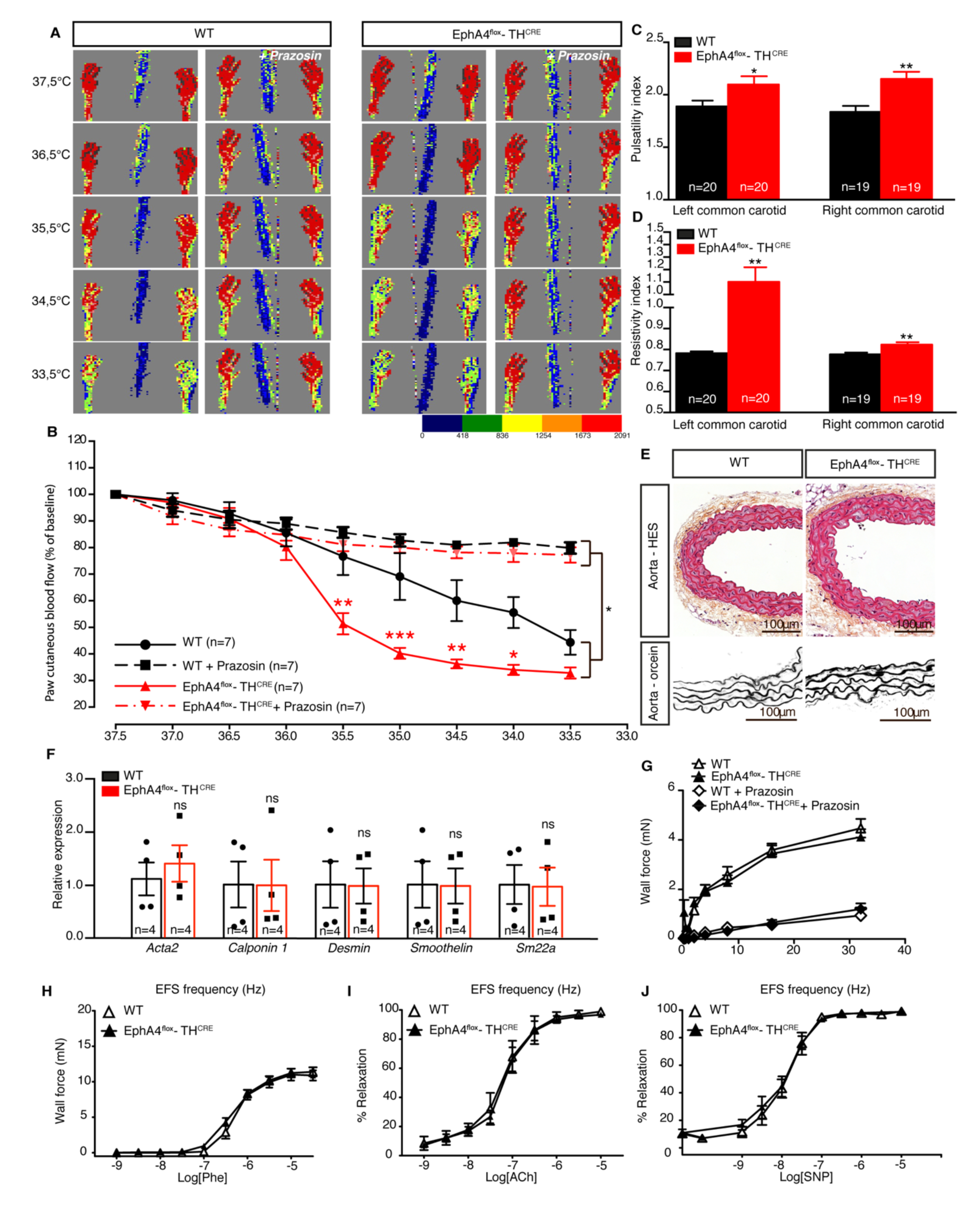
Hyper-innervated arteries show an enhanced vasoconstriction and resistivity but no anatomical, molecular and physiological change. (**A**) Laser Doppler recordings of hind paw cutaneous blood flow of adult WT and EphA4^flox^-TH^CRE^ mice under anesthesia and monitored for body core temperature, without (left) or with injection of prazosin (1mg/kg Intra-peritoneal, right). Red color indicates highest blood flow. (n=7 mice per group). (**B**) Quantification of vasoconstriction in EphA4^flox^-TH^CRE^ mice and WT littermates. For each animal, the measurement of the foot blood flow at 37,5°C was considered as 100%, and the data from other temperatures were expressed as the percentages of the measurement at 37,5°C (% of baseline). (**C and D**) Pulsatility (C) and resistivity (D) indexes of the left and right common carotids of adult WT and EphA4^flox^-TH^CRE^ mice recorded by ultrasound. (**E**) HES (hematoxylin eosin sirius red, top) and orcein (bottom) coloration of transverse sections of aortas from adult WT and EphA4^flox^-TH^CRE^ mice. (**F**) Normalized relative expression of *Acta2*, *Calponin1*, *Desmin*, *Smoothelin*, *Sm22a* mRNA by aortas from adult WT mice and EphA4^flox^-TH^CRE^ littermates. (**G-J**) Isolated rings of mesenteric arteries from adult WT and EphA4^flox^-TH^CRE^ mice were mounted in a wire-myograph and submitted to electrical field stimulation (EFS, **G**). In other arterial rings, contraction induced by phenylephrine (Phe, 1 nmol/L to 30 µmol/L, H) and relaxation induced by acetylcholine (ACh, 1 nmol/L to 10 µmol/L, I) or sodium nitroprusside (SNP, 0.1 nmol/L to 30 µmol/L, J) were measured. ns : not significant; *p<0,05; **p<0.01

Therefore, arterial anatomical and molecular composition as well as physiological response was similar between WT and EphA4^flox^-TH^CRE^ mice. Thus, differences observed in vasoconstriction efficacy and occuring at a higher temperature, as well as enhanced pulsatility and resistivity, are likely associated with arterial sympathetic hyperinnervation. Inhibition of SMC-induced vasoconstriction in presence of Prazosin corroborates the involvement of sympathetic innervation.

We then assessed *ex vivo* vasoreactivity of first (not shown) and second order mesenteric arteries. There was no difference in Phe-mediated vasoconstriction and in Ach and SNP-mediated vasorelaxation between groups (*n*=14 and 18 vessels from 7 and 9 WT and EphA4^flox^-TH^CRE^ mice respectively). Similarly, vasoreactivity was not different between WT and EphA4^flox^-TH^CRE^, and both genotypes responded similarly with or without prazosin (Figure 3, G-J). This observation indicates that arterial wall resistance enhancement in EphA4^flox^-TH^CRE^ is due to the sympathetic innervation and tone, but when *ex vivo* arteries are missing sympathetic input from the sympathetic ganglion chain, arteries of both genotypes behave similarly.

### Sympathetic peripheral arterial resistance induces high blood pressure independently of heart rate and renal regulation

To investigate the potential physiological role of enhanced sympathetic innervation in peripheral resistance arteries, we recorded mice vital parameters including mice activity, arterial blood pressure, heart rate using telemetry (Figure 4A). We recorded parameters during the implant post-surgery recovery, as we assumed this could constitute already a challenging condition. We then let mice recover during 7 days without recording and then recorded freely moving animals for 48 hours. Parameters were normalized over this time window and constituted the basal state of each animal after a full recovery. Finally, we introduced a new adult male in the cage of the recorded animal, inducing an acute stress. We found that whatever the situation, the mean arterial blood pressure (MAP) was significantly and sustainably higher in EphA4^flox^-TH^CRE^ animals compared to WT littermates (Figure 4B). Notably, introduction of a new mate in the cage was sufficient to induce substantial hypertension (Figure 4, A-G). This data provides a proof of concept that an enhanced arterial innervation due to a lack of repulsion of sympathetic axons could result in hypertension in young healthy EphA4^flox^-TH^CRE^ mice. Arterial blood pressure depends on the vascular resistance and the cardiac output. However, heart rate was unchanged in EphA4^flox^- TH^CRE^ animals compared to their WT littermates. Furthermore, heart weight, left ventricle (LV)/total heart, LV ejection fraction and LV shortening fraction as well as 3D- sympathetic innervation (Supplemental Figure 4, A-H) were unaffected suggesting that dysregulation of blood pressure is rather due to the enhanced arterial wall resistivity than to a cardiac effect in our experimental settings.

**Figure 4.**
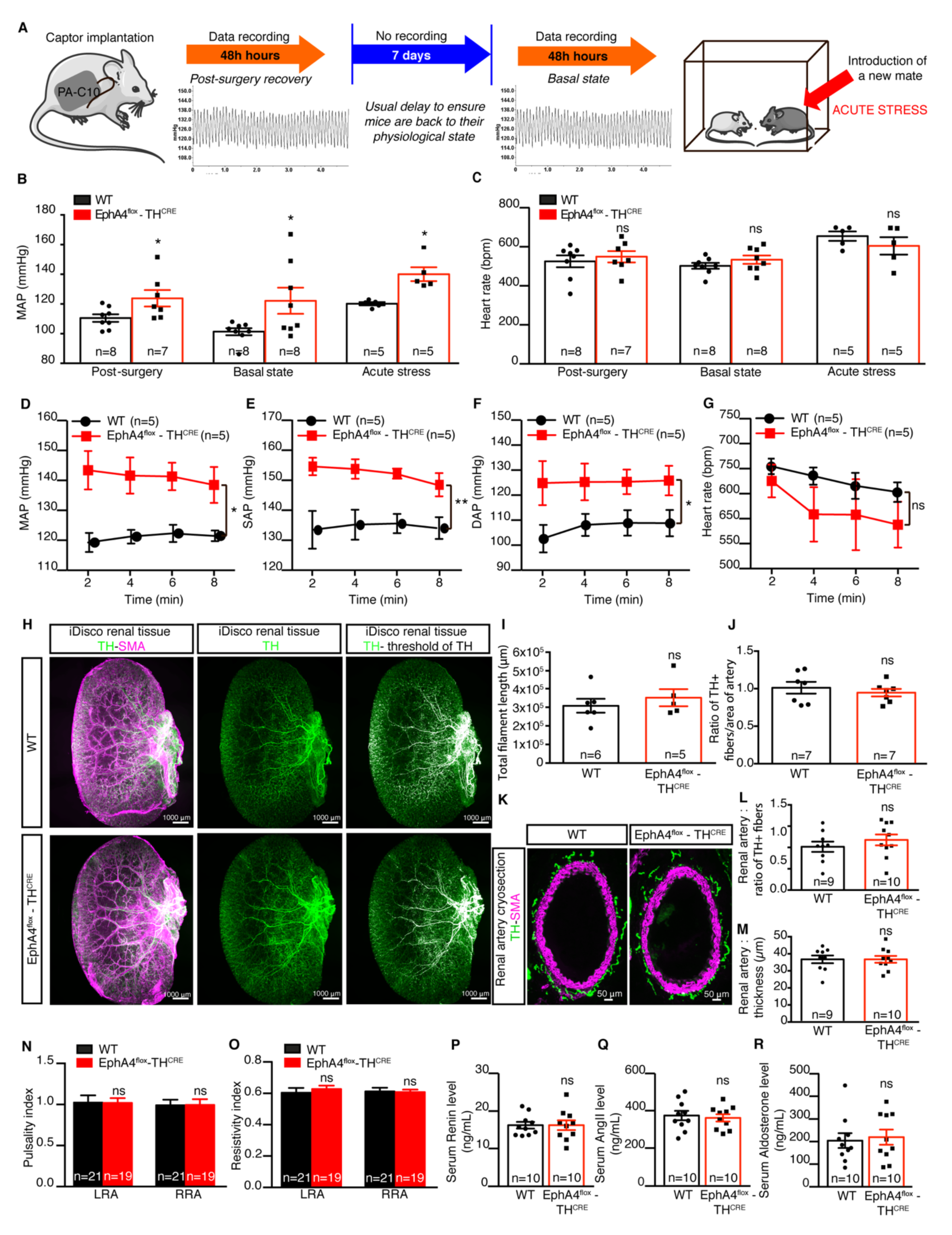
Sympathetic peripheral arterial resistance induces elevated arterial blood pressure independently from heart rate and renal regulations. (**A**) Experimental design of telemetric recordings: after captor implantation, data were recorded during 48 hours post-surgery, followed by 7 days without recording, basal state recording (48 h) and after an acute stress (introduction of an unknown animal within the cage). (**B and C**) Mean arterial pressure (MAP) (B) and heart rate (C) during post-surgery phase, basal state and acute stress phase of adult WT and EphA4^flox^- TH^CRE^ mice. (**D-G**) MAP (D), systolic arterial pressure (SAP, E), diastolic arterial pressure (DAP, F) and heart rate (G) (measurements every 2 minutes during 8 minutes) of adult WT and EphA4^flox^-TH^CRE^ mice during acute stress phase. (**H**) Cleared kidneys (iDisco method) stained for TH (sympathetic nerve fibers, green) and SMC (SMA, magenta) from adult WT and EphA4^flox^-TH^CRE^ mice. The sympathetic nervous network was quantified using TH signal fluorescence intensity (right panels). Large bundles of sympathetic axons appear in white. (**I**) Quantification of the total filament length representing the sympathetic nervous network of kidneys from adult WT and EphA4^flox^-TH^CRE^ mice. (**J**) Quantification of TH+ nerve fibers covering penetrating renal arteries of adult WT and EphA4^flox^-TH^CRE^ mice. (**K**) Immunofluorescent staining of TH (green) and SMA (magenta) on sections of renal main arteries of adult WT and EphA4^flox^-TH^CRE^ mice. (**L and M**) Quantification of the percentage of TH+ nerve fibers covering renal arteries (L) and of arterial wall thickness (M) from adult WT and EphA4^flox^-TH^CRE^ mice. (**N and O**) Pulsatility (N) and resistivity (O) indexes of left and right renal arteries from adult WT and EphA4^flox^-TH^CRE^ mice. (**P-R**) Serum levels of renin (P), angiotensin II (Q) and aldosterone (R) from adult WT and EphA4^flox^-TH^CRE^ mice. ns : not significant; *p<0,05.

EphA4 is expressed within the brain (18). To test a potential central regulation of blood pressure, we identified cells expressing EphA4 within the brain using immunostaining. EphA4-positive cells were distinct from the one expressing TH, so one can reasonably assume that EphA4 was not inactivated within the brain in the EphA4^flox^-TH^CRE^ animals (Supplemental Figure 5, A-E).

*EphA4* global knock-out mice develop kidney malformations, namely hydronephrosis, and thus hypertension (26). We thus investigated general kidney anatomy and hydronephrosis in EphA4^flox^-TH^CRE^ mice. We found no hydronephrosis in those animals, suggesting that hydronephrosis was not due to EphA4 signaling in sympathetic axons but probably due to the role of EphA4 in kidney development and morphogenesis (26). Masson’s trichrome histological analysis revealed no differences in renal tissue and renal arteriole aspects (Supplemental Figure 4I) while renal sympathetic innervation of glomeruli was unaffected. Arterial tree and sympathetic innervation within renal tissue were analyzed in 3D using iDISCO+ protocol. We quantified sympathetic innervation along arteries and within the kidney and found no differences between both genotypes (Figure 4, H-J). Similarly, as renal artery sympathetic innervation could regulate blood pressure and hypertension (27), we quantified renal artery innervation and found no difference (Figure 4, K-M). Hence, EphA4 deletion is not sufficient to induce hyperinnervation, or there is a potential redundancy of guidance cues, at the vSMC or neuronal level, ensuring appropriate rate of innervation within the renal arteries. At the functional level, pulsatility and resistivity index of left and right renal arteries (LRA and RRA, respectively) were not significantly different between both genotype (Figure 4, I and J). Finally, blood pressure is mainly regulated by endocrine function of the kidney and more specifically by the production of Renin, Angiotensin II (AngII) and Aldosterone (28). Serum levels of Renin, AngII and Aldosterone were not different in our experimental conditions (Figure 4, K-M). Therefore, the hypertensive phenotypes in EphA4^flox^-TH^CRE^ mice (Figure 4B) is not due alterations of renal anatomy and function but rather implicates the hyperinnervation-induced enhanced resistivity of peripheral and resistance arteries network.

## Discussion

Interaction between EphrinA4 from vSMC and sympathetic neurons expressing EphA4 mediates axonal repulsion and regulates the level of sympathetic innervation of peripheral arteries. Netrin-1 is also expressed by vSMC to attract sympathetic growth cones toward arteries at the onset of innervation and is involved in the maintenance of arterial innervation (6). We here demonstrated that guidance of arterial innervation is orchestrated as growth cone are first attracted by Netrin-1 secretion via DCC, and then EphrinA4/EphA4 contact-mediated repulsion avoid inappropriate and supernumerary innervation. As sympathetic axons can regenerate, expression of molecules regulating proper innervation rate is needed in adults, allowing the maintenance of appropriate sympathetic innervation. This observation opens the possibility that sympathetic innervation of arteries could be regulated during the entire life, meaning that re-innervation of new organs or new arteries could be possible. This is important in the context of grafts and regenerative medicine, but likely in other fields such as cancer as the level of arterial innervation is altered in some solid tumors (29, 30). On the same note, arterial innervation rate could be a predictive factor for tumor aggressivity and metastasis spreading.

In addition, when treatment of sympathetic denervation is proposed, as for renal artery denervation trials, despite the question of denervation efficacy, the transient status of the denervation process should be taken into account (31).

Regarding the involvement of those guidance molecules in synaptogenesis, while Netrin-1 affects the formation and size of the synapses, EphrinA4 does not seem to regulate synaptogenesis as neither the ratio nor the morphology of synapses is affected. Nevertheless, whether Netrin-1 and Ephrin-A4 are expressed by the same vSMC, or by different cells remain unknown. It has been reported that Netrin-1 and EphrinA4 could potentiate each other effect, probably by involving the same second messenger pathways (32, 33). One can also speculate that some vSMC could be used as “guidepost” to anchor sympathetic innervation. Given that vSMC layers of resistance arteries could be seen as a syncytium and that vSMC are a heterogeneous population, the site of innervation and the level of innervation could thus influence the entire structure. One can therefore speculate that 1) “pacemaker” cells need to have a really refined level of innervation and 2) syncytium receiving an inadequate enhanced innervation would increase the resistivity of the entire muscular tissue layer of the artery. This hypothesis is supported by the observation that EphA4^flox^-TH^CRE^ mice displayed elevated resistivity index observed in EphA4^flox^-TH^CRE^ mice, whereas the general properties and anatomy of the artery (diameter, number of vSMC layer, arterial wall thickness, gene expression) remained unchanged.

Finally, sympathetic innervation of peripheral arteries is increased in EphA4- deleted mice but remains unchanged in the renal artery and the kidney in general. Nevertheless, our model shows an elevated arterial blood pressure in steady-state conditions, which turns into a characteristic hypertension when the wall of the peripheral arterial network is more strongly innervated by sympathetic axons. The increased resistivity of the arterial walls network rises blood pressure, independently of the Renin-Angiotensin-Aldosterone system. We therefore hypothesize that a substantial number of primary and idiopathic hypertension could be of sympathetic origin. Furthermore, high level of arterial sympathetic innervation could contribute or aggravate hypertension in everyday life when patients face stressful or challenging situations. We believe that our results could open up new therapeutic avenues for the treatment of idiopathic hypertension, especially in young subjects and those whose hypertension is resistant to conventional treatments.

## Methods

### Animal study

Experiments used males between P1 and P3 or age-matched littermates. Mice were group housed between 2 and 5 animals per cage with enrichment in a temperature and humidity-controlled animal facility at 22°C under a 12:12-h light: dark circle with free access to standard chow (2018C, Teklad Diets) and water. In telemetry experiments, age-matched male littermates were housed isolated with enrichment in a temperature- and humidity-controlled animal facility at 22°C under a 12:12-h light: dark circle with free access to standard chow (R04, Safe, France) and water.

### Mouse lines

EphrinA4^-/-^, EphA4^-/-^, EphA4^flox^, Ntn1 ^LacZ/+^ and TH^CRE^ mice have been previously described (18, 21, 23, 24, 25). EphA4^flox^ and TH^CRE^ mice have been crossed together to generate EphA4^flox^ - TH^CRE^ mice. EphA4^-/-^ and Ntn1 ^LacZ/+^ mice have been crossed together to generate EphA4^-/-^-Ntn1^LacZ/+^ mice.

### Cell culture

Sympathetic neurons were obtained from Superior Cervical Ganglia (SCG) from post-natal day 1 (P1) pups. Freshly dissected SCG were digested 1h at 37°C in trypsin 1X (Trypsin 10X, Life technologies, diluted 1:10 in DPBS). Sympathetic neurons were dissociated, plated on culture slides (11mm diameter, coated with poly-L-lysine 0,001% (Sigma) and laminin (Sigma) at 10μg/mL) in culture media. Culture media contains 50% Dulbecco’s Modified Eagle Medium (DMEM)-GlutaMAX^TM^ (Gibco®, Life technologies), 50% F12-Nutrient Mixture - GlutaMAX^TM^ (Gibco®, Life technologies), 10% decomplemented Fetal Bovine Serum (Gibco®, Life technologies) 0.2M Penicillin /Streptomycin (Invitrogen) and was supplemented with Nerve Growth Factor (NGF - Sigma) at 20ng/mL. Mesenteric arteries were collected from P15 WT mice and digested at 37°C with elastase (1,25 U/mL, Serabio Technologies) and collagenase (17,5 U/mL, Sigma) for 2 hours. Smooth muscle cells (SMC) were dissociated and plated on culture slides (11mm diameter, coated with poly-L-lysine 0,0 1% and laminin 10μg/mL) in culture media. Culture media contained the same ingredients as the one for sympathetic neurons primary cultures, except for NGF. SMC were maintained in culture for 1 week, changing culture media every two days.

### Binding assay

Rm-EphrinA4/Fc chimera (R&D) was clustered with anti-human Cy^3^ antibody (Sigma) diluted 1: 10 in culture media. Sympathetic neurons were incubated for 5min at 37°C with EphrinA4/Fc 10μg/mL. Controls were anti-human Cy^3^ diluted at 1: 10 in culture media and culture media alone. At the end of the binding experiment, cells were fixed in DPBS/Para-formaldehyde (PFA) 2%/Sucrose 15% 30min at room temperature (RT). Red area per growth cone was imaged with a Leica DMRB Videomicroscope, objective 63X – NA 1.25, equipped with a CCD Coolsnap camera HD monochrome (Photometrics), using the acquisition software Metamorph 7.8 (Molecular Device) and analyzed using ImageJ. Quantification was done 3 or 4 times independently by an observer blinded to the experimental condition.

### Collapse assay

Sympathetic neurons were incubated for 10min at 37°C with a control solution (culture media used for plating) or with a solution of EphrinA4 at the given concentrations. Sympathetic neurons were fixed with DPBS/ paraformaldehyde (PFA) 2%/Sucrose 15% 30min at room temperature (RT) and immunostained with TH. Collapsed or un-collapsed growth cones were scored using standard criteria (34). Quantification was done 3 or 4 times independently by an observer blinded to the experimental condition.

### Cellular immunostaining

Fixed cells were permeabilized with DPBS/0.1% Triton X-100 10min at RT, washed in DPBS, blocked 30min at RT in blocking solution (containing 1% of albumin from Bovine Serum (BSA) – Sigma), incubated in primary antibody diluted in blocking solution 1h at RT, briefly washed in DPBS, incubated in secondary antibody diluted in blocking solution, briefly washed and mounted on slides in mounting medium (Dako). Images were acquired with a Yokogawa CSU-W1 type Spinning Disk, objective 63X (Zeiss PL APO NA 1.4), equipped with Flash 4 Cmos v2+ camera (Hamamatsu), using the acquisition software Metamorph 7.8 (Molecular Device).

### Tissue collection for in situ hybridization

Samples were freshly collected in RNase free conditions, incubated in toluene, rehydrated in bath of decreasing concentration of ethanol, digested with K proteinase 10min at 37°C (Invitrogen) and fixed with DPBS/PFA 4%.

### In situ hybridization

*In situ* hybridization with digoxigenin-labeled mouse Efna4 anti- sense and sense cDNA was performed on whole mesenteric arteries from P2 mice. The bound probes were visualized with alkaline-phosphatase-conjugated fab fragment of antibody to digoxigenin (Boehringer-Mannheim). Images were acquired with a Leica DMRB microscope, objective 10X – PL Fluotar ON 0.30, equipped with a Nikon Camera DXM 1200, using the acquisition software NIS Element (Nikon).

### RNAscope in situ hybridization

*In situ* hybridization was done on sections of mesenteric arteries from P2 pups using the RNAscope® Multiplex Fluorescent Reagent Kit (Advanced Cell Diagnostics, Inc.). *In situ* hybridization protocol was performed as recommended by the manufacturer. Probes against mouse *Efna4* were commercially available form Advanced Cell Diagnostics, Inc.

### Tissue collection for whole-mount immunostaining

Mesenteric arteries from P2 pups and ears from adult mice were collected, conserved in cold DPBS upon dissection and fixed in DPBS/PFA 4% 30min at RT with shaking.

### Whole-mount immunostaining

Samples were blocked 4h in blocking buffer (containing 10% Tris pH=7.4 (Sigma), 0.5% Blocking Reagent (Perkin Elmer), 0.5% Triton X-100, 0.15M NaCl), incubated in primary antibody diluted in blocking buffer over-night (O/N) at 4°C with shaking, washed in washing buffer (containing 10% Tris pH=7.4, 0.05% Triton X-100, 0.15M NaCl), incubated in secondary antibody diluted in blocking buffer 4h at RT with shaking, washed with washing solution and mounted on slides with mounting medium (Dako). Images of mesenteric arteries were acquired with a Leica SP5-MP microscope, objective 63X NA 1.4, using the Leica Acquisition Software 2.4. Images of adult cutaneous arteries were acquired with a Zeiss Axiozoom, objective x260, using the acquisition software Zen. TH+ area per artery was analyzed using ImageJ.

### Tissue collection and cryosection

SCG and mesenteric arteries from adult mice were dissected and prepared for freezing according to previously described protocol (35). Samples were quickly frozen in liquid nitrogen, sectioned in 14μm thick sections and immunostained.

### Section immunostaining

Slides were fixed 10min in cold acetone, drought 30min at RT, blocked in blocking solution (containing 10% Tris pH=7.4 (Sigma), 0.5% Blocking Reagent (Perkin Elmer), 0.5% Triton X-100, 0.15M NaCl) 1h at RT, incubated in primary antibody diluted in blocking solution O/N at 4°C, washed in washing solution (containing 10% Tris pH=7.4, 0.05% Triton X-100, 0.15M NaCl), incubated in secondary antibody diluted in blocking solution 2h at RT, washed in washing solution and covered by coverslips with mounting medium (Dako). TH+, Synaptophysin+ area per artery section were analyzed using ImageJ.

## Antibodies

*Primary antibodies*: anti-ephrinA4 (Abcam) 1/20 on SMC, anti-EphA4 1/100 on sympathetic neurons (Covalab) and on SCG (R&D), anti-Tuj1 1/200 (R&D), anti-tyrosine hydroxylase 1/200 (Millipore), anti-synaptophysin 1/500 (BD Biosciences). *Secondary antibodies*: donkey anti-rabbit 555 and 488 1/200 (Invitrogen), donkey anti-mouse IgG1 488 1/200 (Invitrogen), donkey anti-mouse IgG (H+L) 555 (Invitrogen), Streptavidine Amersham Cy^TM^ (GE Healthcare), donkey anti- goat 555 1/200 (Invitrogen). Coupled antibodies: anti-SMA Cy^3^ 1/200 (Sigma), anti- SMA FITC (Sigma).

### Tissue collection and inclusion for histological studies

Mesenteric arteries were dissected from adult mice, fixed in DPBS/PFA 4% O/N at 4°C with shaking, washed in DPBS, dehydrated in baths of increasing concentration of ethanol, incubated in xylene for several minutes at RT and embedded in paraffin. 14μm-thick sections were stained for elastic fibers using orcein staining. Images were acquired with a Leica DMRB microscope, objective 63X – HCX-PL APO ON 1.40, equipped with a Nikon Camera DXM 1200, using the acquisition software NIS Element (Nikon). After dissection, the kidney, aorta, and heart tissues were fixed for 24 hours in 4% formalin and embedded in paraffin. 4 μm-thick sections were stained with hematoxylin and eosin for all the tissues, Sirius red (HES) (collagen staining) for the aortas and hearts, elastic stain for the aortas, and Masson’s trichrome for the kidneys. Histopathological analysis assessed qualitatively if fibrosis, cellular hypertrophy, arterial wall thickening developed in the mouse target organs. Images were acquired with a Leica DMRB microscope, objective 20X – PL Fluotar ON 0.30, equipped with a Nikon Camera DXM 1200, using the acquisition software NIS Element (Nikon).

Measurement of arterial diameter and thickness of arterial wall were performed on ImageJ.

### Tissue collection and pre-treatment for clarification

Adult mice littermates were sacrificed by cervical elongation. Kidneys and hearts were collected and fixed on PBS/PFA 4% overnight at 4°C and 1h at RT with shaking. Fixed organs were washed in PBS 30 minutes, 3 times at RT with shaking and dehydrated by incubation in baths of increasing concentration of methanol (20%, 40%, 60%, 80% and 100%, 1h each) at RT with shaking. Organs were incubated 1 more hour at RT with shaking and chilled at 4°C before incubation in 66%DCM (dichloromethane)/33%Methanol overnight at RT with shaking. Samples were washed twice in methanol at RT, chilled at 4°C and bleached in 5%H2O2-Methanol overnight at 4°C with shaking. Samples were then rehydrated with methanol series (80%, 60% 40%, 20%, 1 hour each) and washed in PBS-0,2% TritonX-100 (PTx.2) 2x1h at RT with shaking. Samples were incubated in permeabilization solution (20% DMSO, 0,2% Glycine, 80% PTx.2) 2 days at 37°C.

### Immunostaining on whole kidneys and hearts

Pre-treated organs were incubated in blocking solution (84% PTx.2, 0,06% donkey serum, 10% DMSO) 2 days at 37°C with shaking, incubated in primary antibody diluted in PTwH, 5% DMSO, 3% donkey serum, during 10 days at 37°C with shaking, washed in PTwH (PBS 0,2% Tween-20, 0,001% heparin) 1 day at RT with shaking, incubated in secondary antibody diluted in PTwH, 3% donkey serum during 4 days at 37°C with shaking and washed in PTwH for 1 day at RT with shaking.

### Kidneys and hearts clarification

Stained organs were cleared by incubation in 66% DCM/33% Methanol 3 hours at RT with shaking, washed 2 times 15min in methanol 100% and conserved in DBE (di benzyl ether).

### Cell counting on SCG sections

SCG were collected and prepared as described in the « tissue collection and cryosection » paragraph and cut in 30μm thick sections. Number of sympathetic neurons per SCG were estimated by stereology, using the StereoInvestigator Software (BMF Bioscience) coupled to a Nikon Eclipse E800 microscope. Interval between slices: 120μm (1 section out of 4). 8 to 10 slices were counted per SCG. The counting frame of 80μmx80μm and a grid of 150μmx150μm were used. The Coefficient of Error (Gundersen) was inferior or equal to 0.05. The same parameters were used for all the samples.

### Quantitative real time (RT)-qPCR

SCG or mesenteric arteries were collected, conserved in RNAlater (Invitrogen), transferred in lysis buffer from NucleoSpin® RNA XS kit (Macherey-Nagel) and mechanically homogenized in a TissueLyser (Qiagen) for 2 times 2min, at a frequency of 30s^-1^. Total RNA was extracted using the NucleoSpin® RNA XS kit and the concentration was determined by using a Nanodrop 2000c spectrophotometer (Thermo Scientific). First-strand cDNA synthesis was performed by using the Superscript III (Life technologies) in a T100^TM^ Thermal Cycler (BIO-RAD), using 500ng of total RNA input. For qRT-PCR, 5μL of cDNA (1:10 dilution for SCG of P3 and adult EphA4^flox^ – TH^CRE^; 1:5 dilution for SCG of adult EphA4^-/-^ and mesenteric arteries of adult EphA4^flox^ – TH^CRE^) was added to SYBR® Green Jumpstart^TM^ Taq Ready Mix (Sigma). Each sample was analyzed in duplicate and run on a MyiQ^TM^ Single Color Real-Time PCR Detection System (BIO-RAD). Mean dCt values for each target gene were normalized against those of GAPDH and HPRT1 mRNA levels, and corresponding ddCt values were log2-transformed to obtain fold-change values. All primers used in qPCR experiments are Quantitect© primers (Qiagen): Mm_Gapdh_3_SG; Mm_Th_1_SG; Mm_EphA4_1_SG. Others primers were designed by Sigma Aldrich: SMA, sense, GGCATCAATCACTTCAAC, SMA, anti- sense, CTATCTGGTCACCTGTATG; calponin1, sense, AAACAAGAGCGGAGATTTGAGC, anti-sense, TGTCGCAGTGTTCCATGCC; desmin, sense, CGTGACAACCTGATAGAC, anti-sense, TTCTCTGCTTCTTCTCTTAG; smoothelin, sense, CCTCAGATACCTTGGACTC, anti-sense, TTGGCAGGATTTCGTTTC; SM22a, sense, CAACAAGGGTCCATCCTACGG, anti-sense, ATCTGGGCGGCCTACATCA.

### Tissue collection and preparation for Transmission Electronic Microscopy

Mesenteric arteries were collected and conserved in Glutaraldehyde 2% for 24h. Samples were fixed in 2% glutaraldehyde in cacodylate buffer 0.1M pH 7.4 for 2h at 4°C, washed and post-fixed with 1% osmium tetroxide in cacodylate buffer for 1h at 4°C. After an extensive wash (3x10 min) with distilled water they were incubated for 2h in 2% uranyl acetate in water. They were then dehydrated in a graded series of ethanol solutions (2x5min each): 50%, 70%, 80%, 90%, and 100%. Final dehydration was performed twice in 100% acetone for 20 min. Samples were then progressively infiltrated with an epoxy resin, Epon 812® (EMS, Souffelweyersheim, France) : 1 night in 50% resin 50% acetone at 4°C in an airtight container, 2x2h in pure fresh resin at room temperature. They were embedded in the bottom of capsules (Beems® size 3, Oxford Instruments, Saclay, France) and the resin was polymerized at 56°C for 48h in a dry oven. Blocks were cut with an UC7 ultramicrotome (Leica, Leica Microsystemes SAS, Nanterre, France). Semi-thin sections (0.5μm thick) were stained with 1% toluidine blue in 1% borax. Ultra-thin sections (70nm thick) were recovered either on copper (conventional morphology) or nickel (immunoelectron microscopy) grids and contrasted Reynold’s lead citrate (Reynolds, ES (1963). Ultrathin sections were observed with a Hitachi HT7700 electron microscope (Elexience,Verrière-le-Buisson, France) operating at 70 kV. Pictures (2048x2048 pixels) were taken with an AMT41B camera (pixel size: 7.4 μm x7.4 μm). Pictures were processed with the open sources image processing program ImageJ, NIH, Bethesda, USA, when needed.

### Laser doppler experiments

Assessment of cutaneous blood flow was performed with a laser Doppler flowmeter (Moor Instruments, Devon, United Kingdom). Adult mice were anesthetized by inhalation of a mix of oxygen and isoflurane (2% in induction and maintenance, Aerrane, BAXTER, France). Mice were then placed on a heating platform and kept unstressed to reach a body temperature of 37.5°C (monitored by a rectal temperature sensor). Bilateral hind paw skin blood perfusion was assessed. The heating platform was then switched off to let the body temperature of the mice decrease. Each 0.5°C of temperature decreasing, bilateral hind paw skin blood perfusion was assessed. Results are expressed as a percentage of baseline. Same experiment was performed a second time on each animal after the intra-peritoneal injection of Prazosin (Sigma, 1mg/kg).

### Ultrasound investigation

Adult mice were anesthetized with a mix of oxygen and isoflurane (4% induction, 1.5-2% maintenance). All acquisitions were done using a 40Hz MS-550S MicroScan^TM^ array transducer (Visualsonics, Inc) connected to Vevo2100 FUJIFILM Visualsonics, Inc (VisualSonics, Toronto, Ontario, Canada). Parasternal Long Axis (PSLA), Short Axis (SAX) and arterial views were acquired in B-mode to assess anatomical aspect of the heart and different arteries. SAX and arterial views were also acquired in M-mode to measure directly on the Vevo2100 the thickness of the left ventricular walls, the diameter and the thickness of carotids and renal arteries. Ejection Fraction (EF) was calculated using the formula 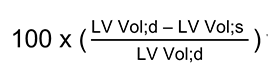 where LV Vol;d stands for “Left Ventricle Volume in diastole” LV Vol;d and LV Vol;s stands for Left Ventricle Volume in systole. LV Vol;d was calculated using the formula 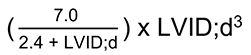 x LVID;d^3^ where LVID;d stands for “Left Ventricular Internal Diameter in diastole” and LVID;s stands for “Left Ventricular Internal Diameter in systole”. LVID;d and LVID;s were measured on M-mode acquisitions. Fractional Shortening (FS) was calculated using the formula 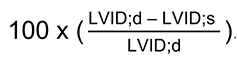. Weight of the LVID;d Left ventricle was calculated on the basis of M-mode measurements, using the formula: (1.053 x ((LVID;d + LVPW;d + IVS;d)^3^ – LVID;d^3^)) x 0.8 where LVPW;d is the thickness of the anterior wall of the left ventricle in diastole and IVS;d is the thickness of the inter- ventricular septum in diastole. Acquisitions in pulse-waved doppler (PW-mode) of carotids and renal arteries were also done to assess blood flow in these arteries. All measurements were done between respiratory movements to avoid bias. Pulsatility index of carotids and renal arteries were calculated using the formula 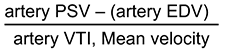 where PSV stands for “Peak Systolic Velocity”, EDV stands for “End Diastolic Velocity” and VTI stands for “Velocity Time Integral”. PSV, EDV and VTI were measured on PW-mode acquisitions. Resistivity index of carotids and renal arteries were calculated using the formula 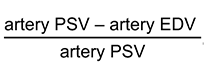.

### Pharmacological profile of isolated mesenteric arteries

Segments of mesenteric arteries were mounted in a wire-myograph (Danish Myo Technology, Denmark) as previously described (36). Electrical field stimulation (EFS) was applied by means of two platinum electrodes placed on either side of the rings and connected to a stimulator (S-900 Stimulator Cornerstone by Dagan). Stimulation was held at 0.02–32 Hz frequency with amplitude of 12 V and pulse duration of 1 ms delivered as 1 min trains (frequency 0.2 Hz, 0.5 Hz, 1Hz, 2Hz, 4Hz, 8Hz, 16Hz and 32Hz) with 10 minutes between 2 stimulations. In other arterial segments, cumulative concentration-response curves (CRCs) to phenylephrine (1 nmol/L to 30 µmol/L) was performed. CRCs to acetylcholine (ACh, 1 nmol/L to 10 µmol/L) or sodium nitroprusside (SNP, 0.1 nmol/L to 30 µmol/L) was obtained after precontraction with phenylephrine (1 µmol/L). Surgery Mice were anesthetized initially with 5% isoflurane in an oxygen stream and maintained on 2-3% isoflurane. To reduce pain, mice received 2 injections of meloxicam (2mg/kg per injection) at 24h interval. Mice were kept on a heating pad throughout implantation of the BP telemeter (TA11PA-C10, Data Science International, St. Paul, MN). The catheter was inserted into the left common artery. This method has been previously described (37). The telemetric transmitter probe was positioned subcutaneously on the right flank. After the mice had recovered from the anesthesia in a warm (37°c) box, they were housed in individual cages placed on top of the telemetric receivers in a light-dark cycled recording room.

### Blood pressure measurements

Study of blood pressure by telemetry was performed on separate, dedicated sets of mice to avoid interference by other measurements. Blood pressure, heart rate (HR), and locomotor activity were monitored 48hours following the surgical implantation of telemetric transmitter, and the 8th and 9th day after the surgery by telemetric recording in conscious, freely moving animals as previously described (37, 38), every hour during 2 minutes. Radiotelemetry probes (model TA11PA-C10; Data Science International, St. Paul, MN) were implanted in age-matched adult mice (2 to 4 months old). Data were analyzed using the Dataquest ART analysis software. Introduction of an unknown male mate was used as an acute stressful stimulus. Activity, blood pressure and heart rate were continuously recorded during this stressful stimulus.

### Serum sampling

200µL of blood were sampled per mice twice at 24h interval at facial vein, using specifically designed lancets (Bioseb). Blood samples were let at RT 30min to coagulate with heparin 10% and centrifugated 10min at 4°C 2000g. Serum were sampled and kept at -80°C.

### ELISA

Serum levels of renin (ThermoScientific), angiotensin-II (EnzoLife Science) and aldosterone (EnzoLife Science) were assessed by ELISA dosages. Serums were gently thawed on ice and diluted at 1/10 (renin and angiotensin) or 1/20 (aldosterone) in respective assay buffers. ELISA dosages were performed according to manufacturer’s instructions. Regression curves were performed and final concentrations calculated using GraphPad Prism 6 software.

### Statistics

For all statistical analyses, GraphPad Prism 6 software was used. All replicate numbers (number of mice analyzed, unless otherwise indicated) are indicated in the figures. Only male mice were included in analyses to avoid impact of sexual cycle on our studies. No statistical methods were used to pre-determine sample size. When possible, all analysis were done blind to genotype and/or treatment groups. Error bars represent SEM in all figures. *P* values of less than 0.05 were considered significant in all experiments. All tests performed were two-tailed. For comparison between two groups, a non-parametric Mann-Whitney t-test was used. For assessment between more than two groups, one-way ANOVA with multiple comparisons (Dunn’s test) was used and for assessment between two independent variables, two-way ANOVA with multiple comparisons (Bonferroni’s test) was used. In telemetry studies (acute stress), two-way ANOVA was used to assess differences between genotypes.

### Study approval

Animal experiments were performed according to ethical regulations and approved protocols by the CIRB ethical committee (n°005) and the CEF ethical committee (n°59) under the Apafis agreement number 6951.

## Author contributions

I.B. and E.S. conceived the study. S.M., E.S., J.V. and E.V. conducted experiments. E.S., S.M., E.V., J.V., V. M., and I.B. analyzed data. I.B. wrote the manuscript. All authors reviewed and edited the manuscript. E.S. and S.M. have contributed equally to the work and share co-first authorship. They are listed on this order as E.S. initiated the study and generated the first results whereas S.M. joined the study later. E.S. and S.M. agree on this assignment of authorships.

## Acknowledgements

We would like to thank Guy Malkinson for critical reading of the manuscript, and Bing- Cheng Wang for providing EphrinAs TKO mice tissues. This work was supported by grants from Inserm, Agence Nationale de la Recherche (ANR NIRVANA) and Fondation pour la Recherche Médicale (FRM). We gratefully acknowledge support by Ligue contre le Cancer and MemoLife Labex (ST), FRM (ES).

Address correspondence to : Isabelle Brunet, Center for Interdisciplinary Research in Biology (CIRB), College de France, CNRS, INSERM, Université PSL, 75005 Paris, France. Phone: + 33 144271693 ;

---

**Supplemental Figure 1.**
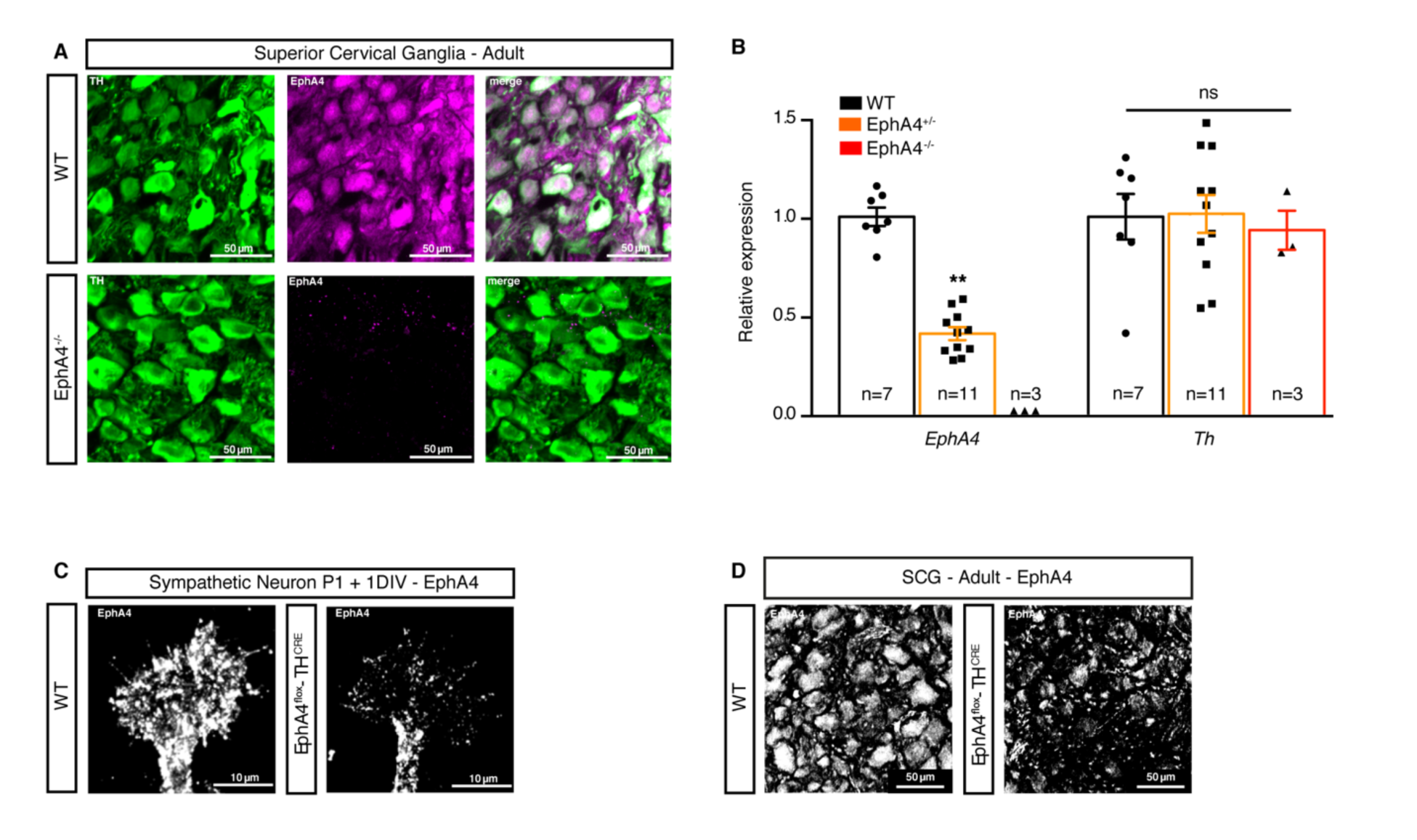
Neuronal loss of EphA4 expression in full knock-out and TH-specific mice. (**A**) Immunofluorescent staining on transverse section of SCG from adult WT and EphA4^-/-^ littermates. Neurons from WT express TH (green) and EphA4 (magenta), whereas EphA4 staining is lost in EphA4 ^-/-^. (**B**) Relative normalized expression of *Th* and *Epha4* mRNA in SCG from adult WT and EphA4^flox^-TH^CRE^. (**C**) Immunofluorescent staining of sympathetic axons and growth cone from WT and EphA4^flox^-TH^CRE^ P1 mice, cultured 1 day *in vitro*. Neuronal EphA4 expression appears in white. (**D**) Immunofluorescent staining of a transverse section of SCG from adult WT and EphA4^flox^-TH^CRE^ mice. EphA4 expression appears in white. ** p<0.01.

**Supplemental Figure 2.**
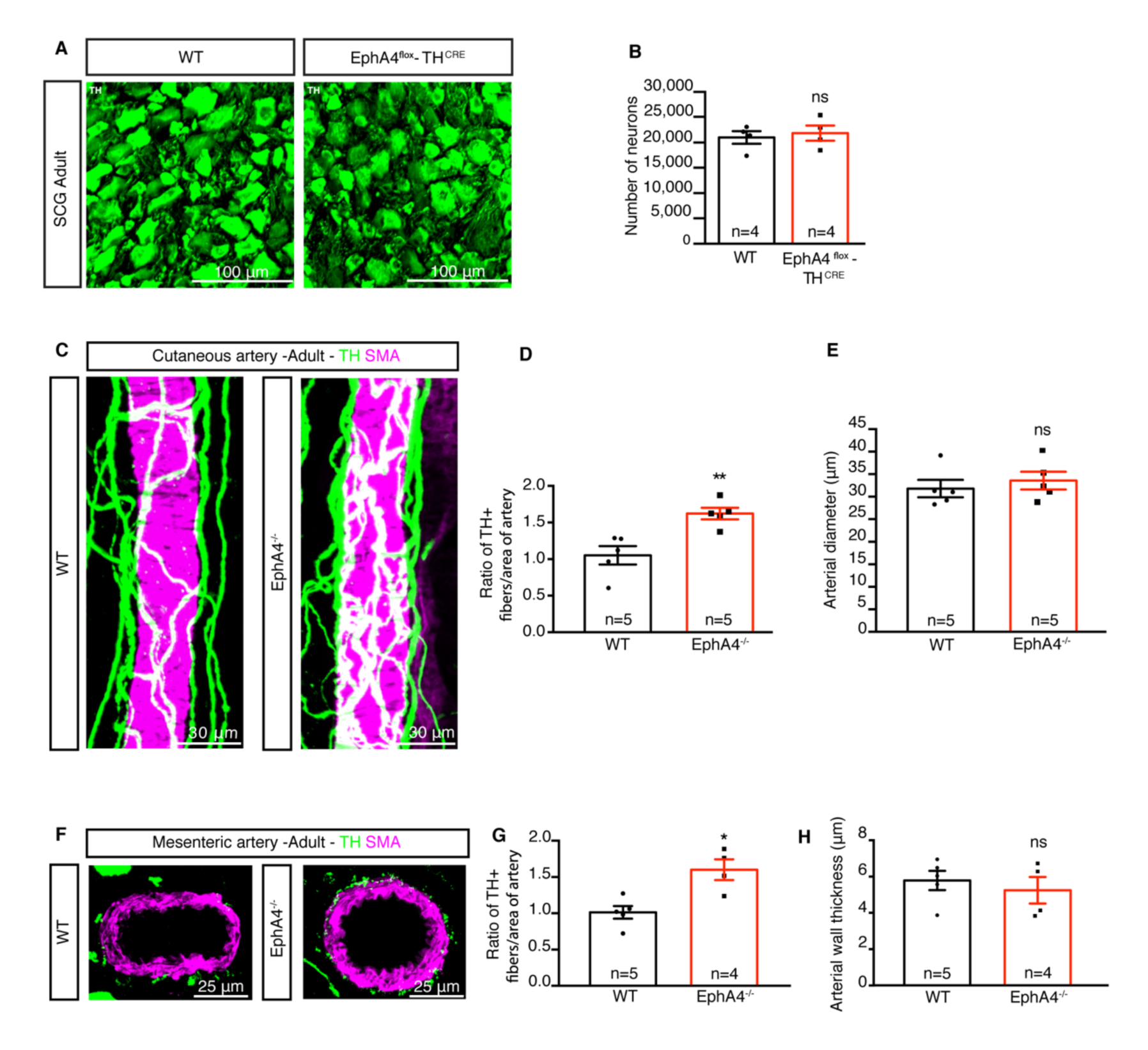
Enhanced arterial innervation in genetically inactivated EphA4 adult mice, while number of neurons per sympathetic ganglia remain unchanged. (**A**) Immunofluorescent staining of a transverse section of SCG from adult WT and EphA4^flox^-TH^CRE^ littermate. Neurons express TH (green). (**B**) Number of TH+ neurons in an SCG from adult WT and EphA4^flox^-TH^CRE^ mice. (**C**) Whole-mount immunofluorescent staining of cutaneous arteries (ears) from adult WT (left) and EphA4 ^-/-^mice (right). Sympathetic nerves expressing TH appear in green whereas smooth muscle cells expressing SMA are shown in magenta. (**D**) Quantification of TH+ nerve fibers covering cutaneous arteries from adult WT and EphA4 ^-/-^mice. (**E**) Quantification of the diameter of cutaneous arteries from adult WT and EphA4 ^-/-^ mice. (**F**) Immunofluorescent staining of sections of mesenteric arteries from adult WT and EphA4 ^-/-^mice stained for TH (green), SMA (magenta). (**G**) Quantification of TH+ nerve fibers on mesenteric arteries from adult WT and EphA4 ^-/-^mice. (**H**) Quantification of the diameter of mesenteric arteries from adult WT and EphA4 ^-/-^mice. ns: not significant; *p<0.05; **p<0.01.

**Supplemental Figure 3.**
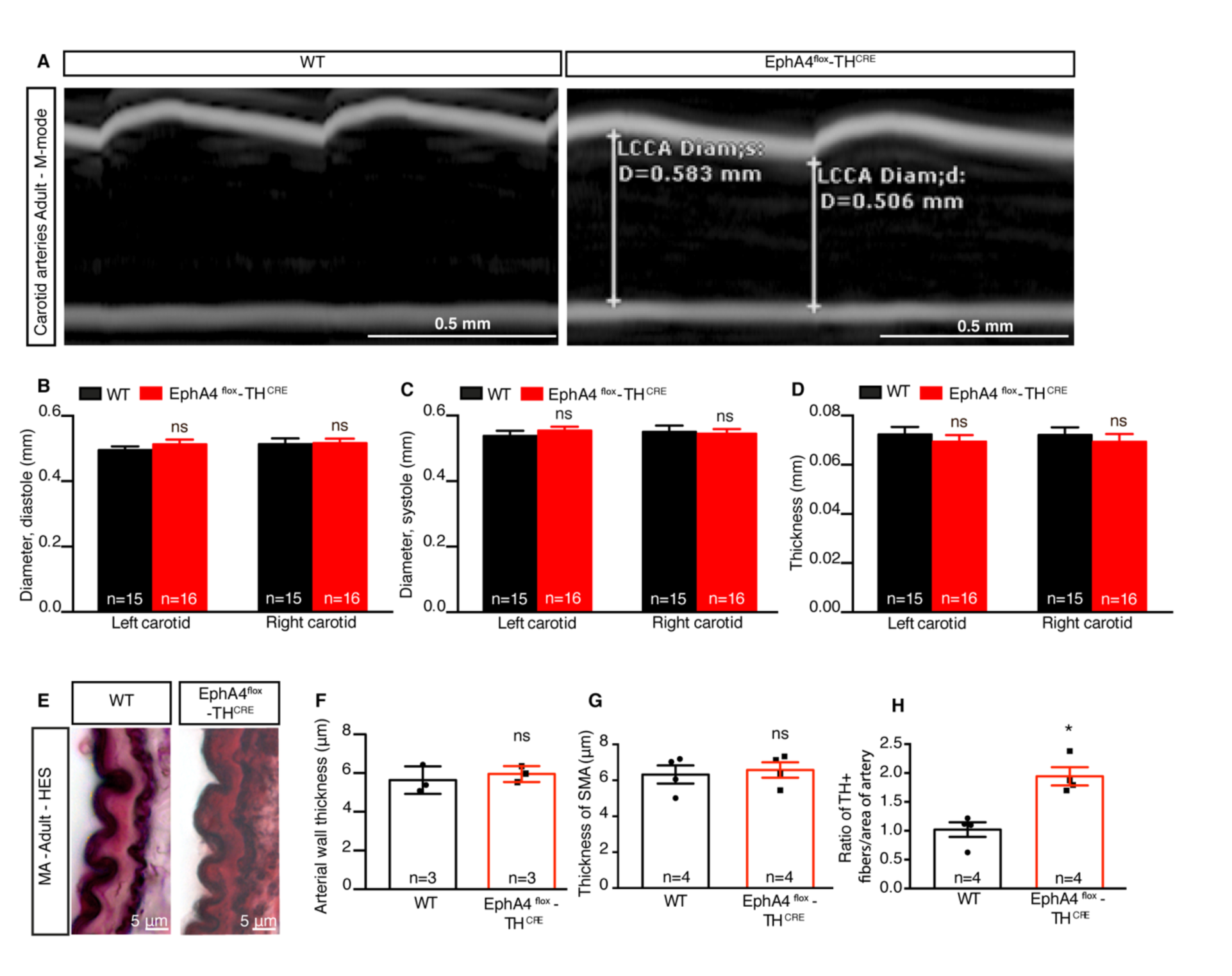
Arterial wall properties with normal or enhanced sympathetic innervation. (**A**) M-mode representative images of the Left Common Carotid Arteries (LCCA) from adult WT and EphA4^flox^-TH^CRE^ mice. Diameter was quantified during diastole and systole phases. (B and C) Quantification of carotid diameter during diastole (**B**) and during systole (**C**) of the left and right common carotids of adult WT and EphA4^flox^- TH^CRE^ mice. (**D**) Quantification of thickness of left and right common carotids from adult WT and EphA4^flox^-TH^CRE^ littermates. (**E**) Transverse sections of mesenteric arteries from adult WT and EphA4^flox^-TH^CRE^ littermate, colored with HES (hematoxylin eosin and sirius red). (**F**) Quantification of the arterial wall thickness of mesenteric arteries from adult WT and EphA4^flox^-TH^CRE^ mice. (**G**) Quantification of the arterial wall thickness of mesenteric arteries from adult WT and EphA4^flox^-TH^CRE^ mice (immunofluorescent staining of SMA was used). (**H**) Quantification of TH+ nerve fibers covering mesenteric arteries from sections quantified in (F and G). ns: not significant; *p<0.05.

**Supplemental Figure 4.**
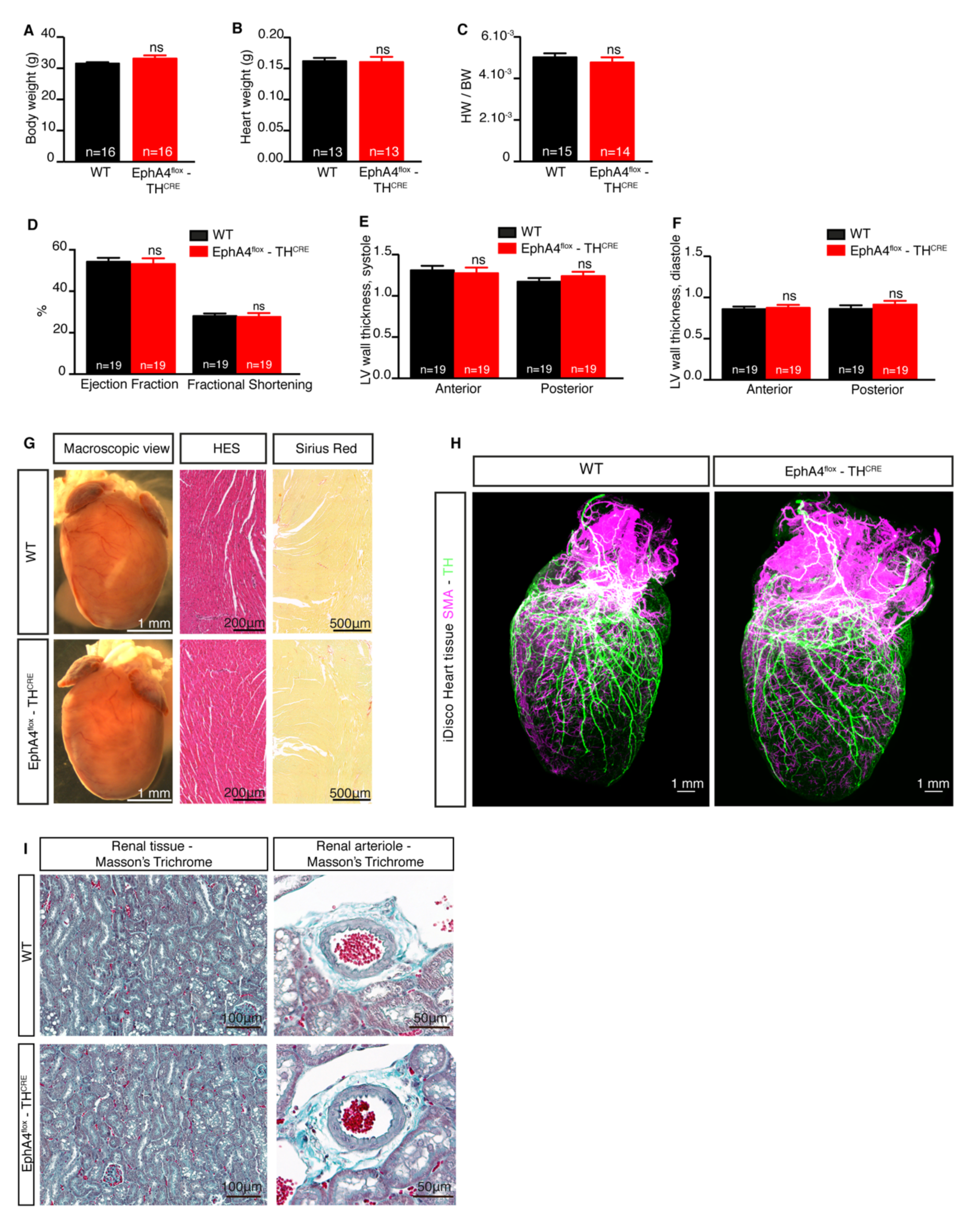
Characterization of EphA4^flox^-TH^CRE^ mice body weight, heart and kidney. (**A**) Body weight of adult WT and EphA4^flox^-TH^CRE^ littermates. (**B**) Heart weight of adult WT and EphA4^flox^-TH^CRE^ mice. (**C**) Ratio of heart weight to body weight of adult WT and EphA4^flox^-TH^CRE^ mice. (**D**) Quantification of the percentage of ejection fraction and fractional shortening of hearts from adult WT and EphA4^flox^-TH^CRE^ mice. (**E and F**) Anterior and posterior left ventricular wall thickness during systole (E) and diastole (F) from adult WT and EphA4^flox^-TH^CRE^ mice. (**G**) Macroscopic view (left panel) and transverse sections of heart from adult WT and EphA4^flox^-TH^CRE^ mice, colored with HES (hematoxylin, eosin and sirius red, middle) and Sirius Red alone (right). (**H**) Snapshots of a 3D view of cleared hearts from adult WT and EphA4^flox^-TH^CRE^ mice (Imaris software). Nerve fibers expressing TH are marked in green, arteries labeled with SMA appear in magenta. (**I**) Transverse sections of kidneys from adult WT and EphA4^flox^-TH^CRE^ mice, colored with Masson’s Trichome, renal tissue (left) and close-up view of renal arteriole (right). ns: not significant

**Supplemental Figure 5.**
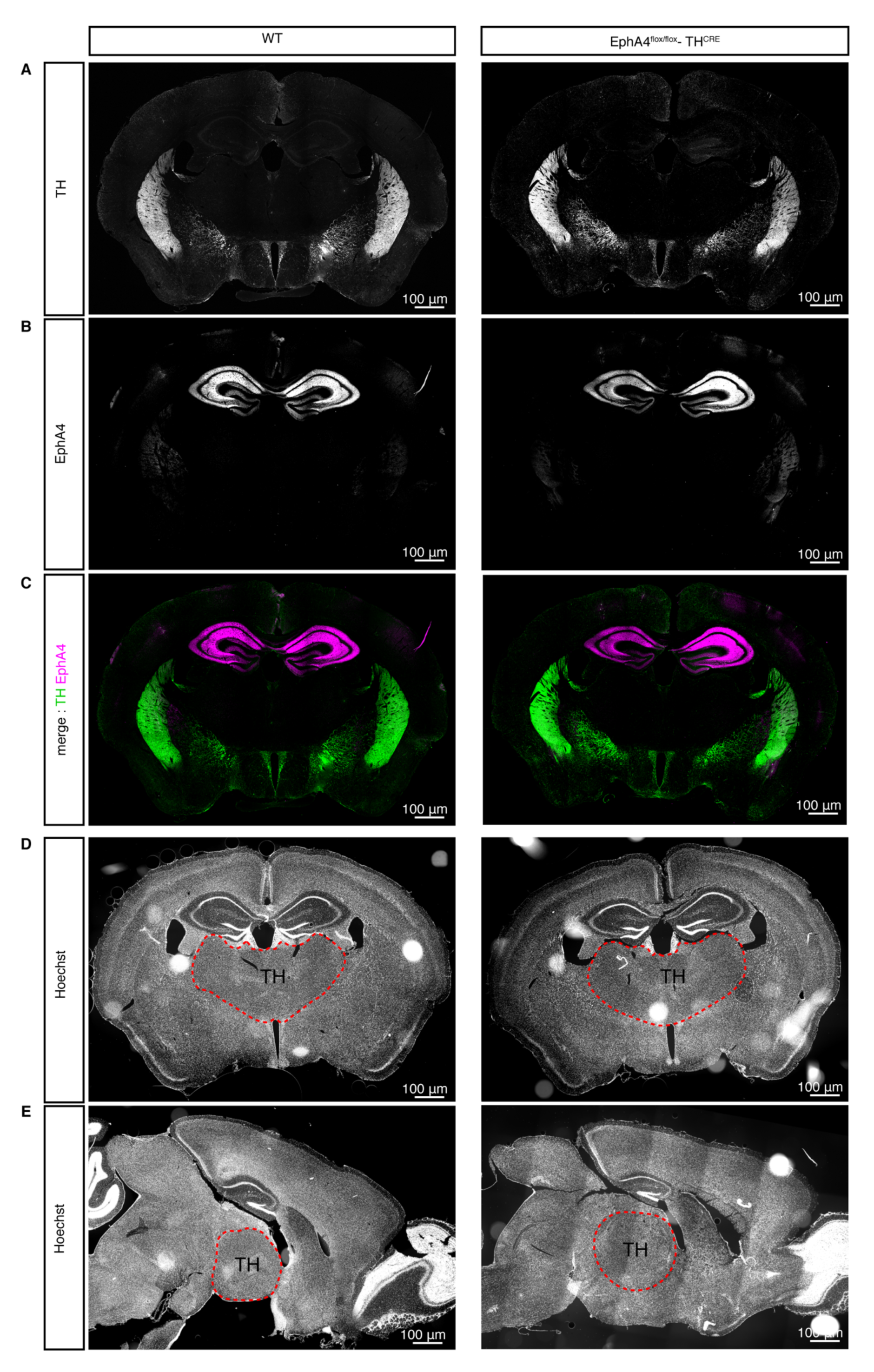
Central Nervous System (CNS) expression of TH and EPHA4 in WT and EphA4^flox^-TH^CRE^ mice. (**A-C**) Immunofluorescent staining of an adult brain coronal section of WT and EphA4^flox^-TH^CRE^ mice. Neurons express TH (green) and EphA4 (magenta). Note no colocalization of EphA4 and TH staining. (**D and E**) Immunofluorescent staining of a coronal (**D**) and sagittal (**E**) sections of a brain from an adult WT and EphA4^flox^-TH^CRE^ mice. Nucleus are stained with Hoechst.

